# Enhancer connectome in primary human cells reveals target genes of disease-associated DNA elements

**DOI:** 10.1101/178269

**Authors:** Maxwell R. Mumbach, Ansuman T. Satpathy, Evan A. Boyle, Chao Dai, Benjamin G. Gowen, Seung Woo Cho, Michelle L. Nguyen, Adam J. Rubin, Jeffrey M. Granja, Katelynn R. Kazane, Yuning Wei, Trieu Nguyen, Peyton G. Greenside, M. Ryan Corces, Josh Tycko, Dimitre R. Simeonov, Nabeela Suliman, Rui Li, Jin Xu, Ryan A. Flynn, Anshul Kundaje, Paul A. Khavari, Alexander Marson, Jacob E. Corn, Thomas Quertermous, William J. Greenleaf, Howard Y. Chang

**Affiliations:** Center for Personal Dynamic Regulomes, Stanford University School of Medicine, Stanford, CA 94305, USA; Department of Genetics, Stanford University School of Medicine, Stanford, CA 94305, USA; Department of Pathology, Stanford University School of Medicine, Stanford, CA 94305, USA; Department of Molecular and Cell Biology, University of California, Berkeley, CA 94720, USA; Innovative Genomics Institute, University of California, Berkeley, CA, 94720, USA; Department of Microbiology and Immunology, University of California, San Francisco, CA 94143, USA; Program in Epithelial Biology, Stanford University School of Medicine, Stanford, CA, 94305, USA.; Chan Zuckerberg Biohub, San Francisco, CA 94158, USA; Department of Medicine, Division of Cardiovascular Medicine, Stanford University School of Medicine, Stanford, California 94305, USA.; Biomedical Sciences Graduate Program, University of California, San Francisco, CA 94143, USA; Department of Applied Physics, Stanford University, Stanford, CA, 94305, USA

**Author notes:** These authors contributed equally (Maxwell R. Mumbach, Ansuman T. Satpathy, Evan A. Boyle, Chao Dai). These authors jointly directed this work (William J. Greenleaf, Howard Y. Chang).

## Abstract

The challenge of linking intergenic mutations to target genes has limited molecular understanding of human diseases. Here, we show that H3K27ac HiChIP generates high-resolution contact maps of active enhancers and target genes in rare primary human T cell subtypes and coronary artery smooth muscle cells. Differentiation of naïve T cells to T helper 17 cells or regulatory T cells creates subtype-specific enhancer-promoter interactions, specifically at regions of shared DNA accessibility. These data provide a principled means of assigning molecular functions to autoimmune and cardiovascular disease risk variants, linking hundreds of noncoding variants to putative gene targets. Target genes identified with HiChIP are further supported by CRISPR interference and activation at linked enhancers, by the presence of expression quantitative trait loci, and by allele-specific enhancer loops in patient-derived primary cells. The majority of disease-associated enhancers contact genes beyond the nearest gene in the linear genome, leading to a four-fold increase of potential target genes for autoimmune and cardiovascular diseases.

---

Gene expression programs are intimately linked to the hierarchical organization of the genome. In mammalian cells, each chromosome is organized into hundreds of megabase-sized topologically associated domains (TADs), which are conserved from early stem cells to differentiated cell types^1^. Within this invariant TAD scaffold, cell type-specific enhancer-promoter (E-P) interactions establish regulatory gene expression programs^2^. Standard methods require tens of millions of cells to obtain high-resolution interaction maps and confidently assign E-P contacts^3-5^. Thus, the principles that govern E-P conformation in disease-relevant patient samples are incompletely understood. This gap in understanding is particularly problematic for interpreting the molecular functions of inherited risk factors for common human diseases, which reside in intergenic enhancers or other non-coding DNA features in up to 90% of cases^6-9^. Such disease-relevant enhancers may not influence the expression of the nearest gene (often reported as the default target in the literature), and instead act in a cell-type specific manner on distant target genes residing up to hundreds of kilobases (kb) away^2,10-14^. Recently, systematic perturbations of regulatory elements in select gene loci have shown that effects of individual regulatory elements on gene activity can be predicted from the combination of (i) enhancer activity [marked by histone H3 lysine 27 acetylation (H3K27ac) level] and (ii) enhancer-target looping^5,15^. Here we leverage this insight to capture the combination of these two types of information genome-wide in a single assay, mapping the enhancer connectome in disease-relevant primary human cells.

## RESULTS

### H3K27ac HiChIP identifies functional enhancer interactions

We recently developed HiChIP, a method for sensitive and efficient analysis of protein-centric chromosome conformation^16^. Cohesin HiChIP in GM12878 cells identified similar numbers of loops as *in situ* Hi-C (~10,000) with high correlation (R = 0.83), demonstrating that HiChIP captures loops with high sensitivity and specificity. Here, we evaluated the enhancer and promoter-associated mark H3K27ac^17-19^ as a candidate factor to selectively interrogate E-P interactions genome-wide. We performed H3K27ac HiChIP in mouse embryonic stem (mES) cells to compare to cohesin HiChIP (**Supplementary Fig. 1a, Supplementary Table 1**)^16^. 3,552 of 4,191 H3K27ac HiChIP loops in mES cells were also identified by cohesin HiChIP. The H3K27ac-biased loops (log_2_ fold-change > 1) spanned shorter distances than cohesin-biased loops, and were enriched for H3K27ac ChIP-seq peaks (78.9%; **Supplementary Fig. 1b-f, Supplementary Table 2**). Moreover, systematic titration of input material showed H3K27ac HiChIP retained high signal fidelity and reproducibility from 25 million to 50,000 cells as input material (loop signal correlation = 0.918; **Supplementary Figs. 2 and 3**). Therefore, H3K27ac HiChIP identifies high-confidence chromatin loops focused around enhancer interactions from limited cell numbers.

In order to capture (i) conformational change during T cell differentiation and (ii) cell type-specific chromatin contacts of autoimmune risk variants in protective and pathogenic T cell types, we performed H3K27ac HiChIP on primary human Naïve T cells (CD4+CD45RA+CD25^-^CD127^hi^), regulatory T cells (T_re_g; CD4+CD25+CD127^low^) and T helper 17 cells (T_H_17; CD4+CD45RA^-^CD25^-^CD127^hi^CCR6+CXCR5^-^) directly isolated from donors (Fig. la,b **and Supplementary Fig. 4a**)^20,21^. T_H_17 cells were sorted to include autoimmune disease-relevant pathogenic TH17 cells and to exclude follicular helper T cells with a distinct surface phenotype and immune function (**Supplementary Fig. 4a**)^22-24^. Peripheral blood CD4+ T cells were isolated from three healthy subjects, isolated by FACS, and subjected to H3K27ac HiChIP. HiChIP libraries from each subset were high quality; greater than 40% of the reads represented unique paired-end tags (PETs) (**Supplementary Fig. 4b-d and Supplementary Table 1**). Furthermore, libraries exhibited high 1D signal enrichment at enhancers and promoters, and globally recapitulated publically available H3K27ac ChIP-seq datasets (74.7% overlap of ChIP-seq and 1D HiChIP peaks; Fig. 1c)^25^. Inspection of the interaction matrix at progressively higher resolution revealed chromatin compartments, TADs, and focal loops, as previously reported in high-resolution Hi-C and HiChIP analyses from cell lines (Fig. 1b)^4,16^. Importantly, H3K27ac HiChIP maps were capable of identifying focal interactions at 1 kb resolution, which is comparable to *in situ* Hi-C maps generated from 100-fold more cells and sequenced to 13-fold greater depth^4^ (Fig. 1b).

**Fig. 1.**
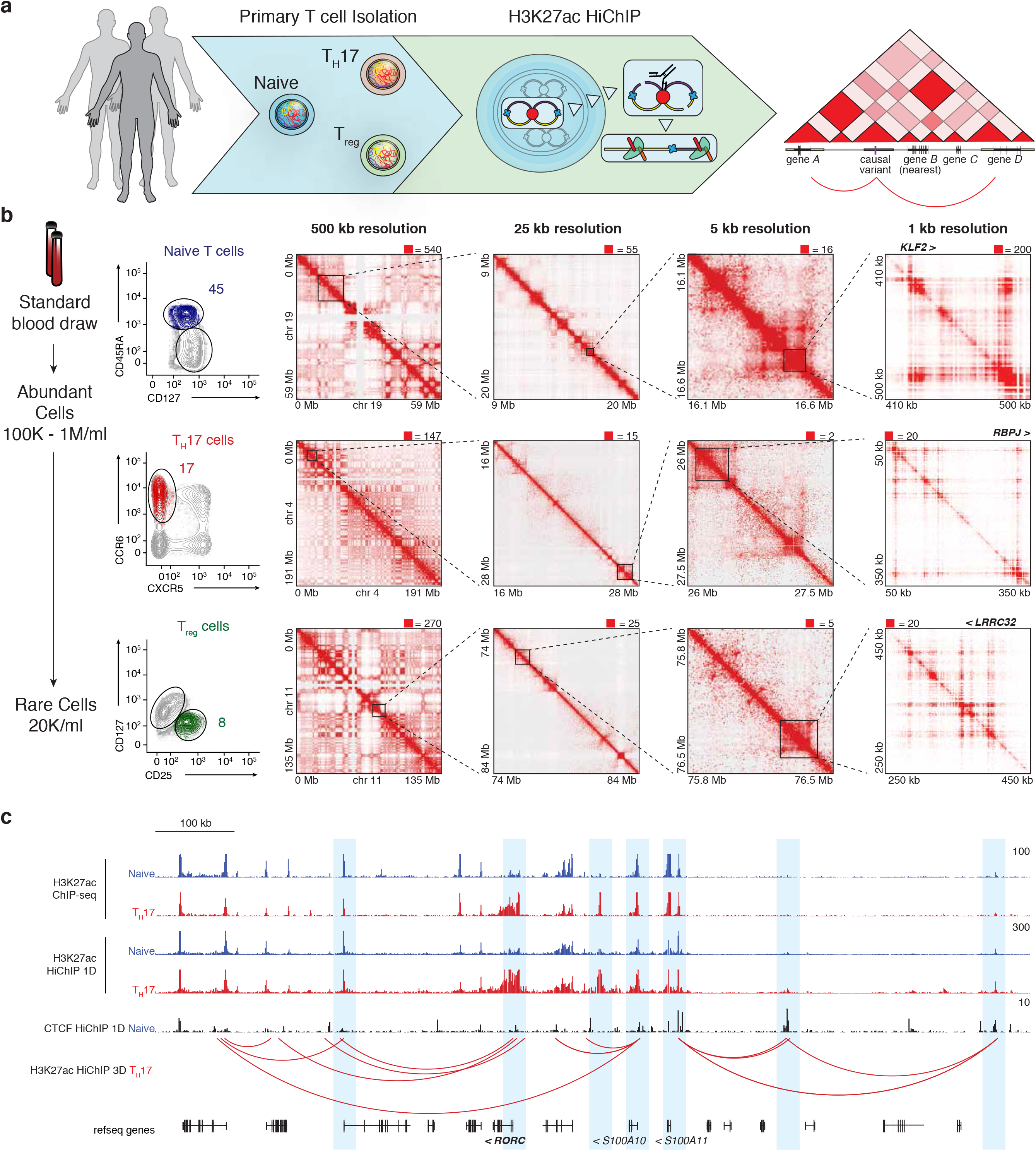
HiChIP identifies high-resolution chromosome conformation in primary human T cells. **(A)** Primary T cell H3K27ac HiChIP experimental outline. **(B)** (Left) FACS strategy for Naïve, Th17, and Treg cells from total peripheral blood CD4+ T cells. Number represents percent of total CD4+ T cells within that gate. (Right) Knight-Ruiz (KR) matrix-balanced interaction maps for Naïve, Th17, and Treg cells at 500 kb, 25 kb, and 5 kb resolution, and raw interaction maps at 1 kb resolution, centered on the *KLF2, RBPJ*, and *LRRC32* loci. **(C)** HiChIP 1D and 3D signal enrichment at the *RORC* locus in TH17 over Naïve T cells.

Previous saturation perturbation screens demonstrated that functional enhancers can be identified by integrating H3K27ac ChIP-seq signal with chromosome conformation contact strength (Hi-C)^5^. Since H3K27ac HiChIP combines these two components into one assay, we reasoned that HiChIP signal, which we term Enhancer Interaction Signal (EIS), should identify functional regulatory elements. To validate this prediction, we first generated H3K27ac HiChIP maps in a chronic myelogenous leukemia cell line (K562) as a direct comparison to published high-resolution CRISPR interference (CRISPRi) screens^5^. We then examined the 3D enhancer landscape of the *MYC* and *GATA1* loci using virtual 4C (v4C) analysis, where a specific genomic position is set as an anchor viewpoint, and all interactions occurring with that anchor are visualized in 2D^16^. v4C analysis of the *MYC* promoter demonstrated that EIS in K562 cells captured all functional enhancers identified in the CRISPRi screen (Fig. 2a). Analysis of the *GATA1* locus demonstrated a similar agreement between both methods (Fig. 2b). Quantitatively, EIS in K562 cells was significantly correlated with CRISPRi score in the same cell type, whereas EIS in GM12878 (GM; B cell lymphoblast) cells was not correlated with K562 CRISPRi (Spearman’s rho = 0.332 and 0.145; p-value = 9.25 x 10^-5^ and 0.1246; Fig. 2c).

**Fig. 2.**
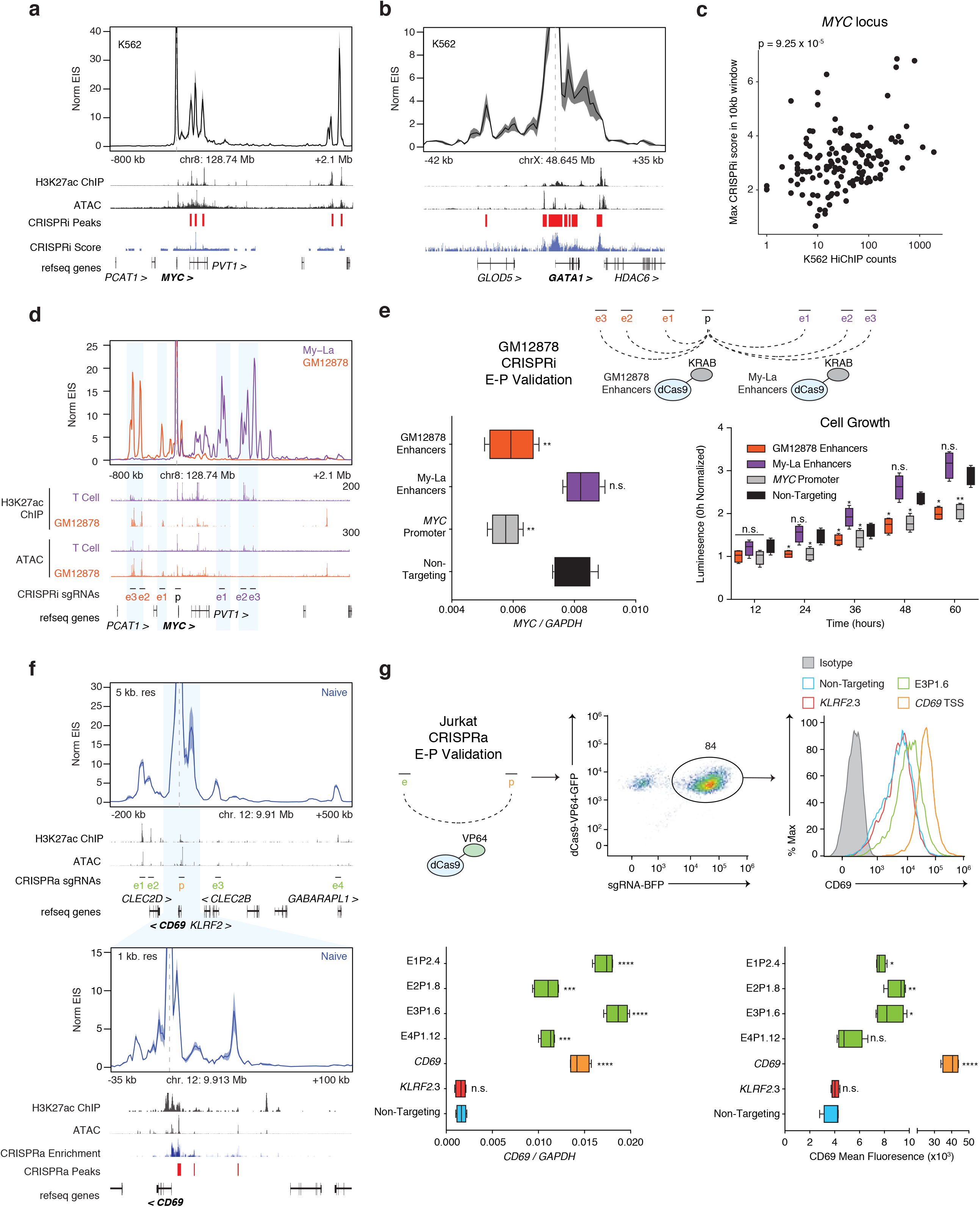
Validation of regulatory elements identified by H3K27ac HiChIP with CRISPR interference and activation. **(A)** Interaction profile of the *MYC* promoter in K562 H3K27ac HiChIP at 10 kb resolution. K562 H3K27ac ChIP-Seq is from ENCODE. CRISPRi-validated regulatory regions in K562 cells are indicated^5^. **(B)** Interaction profile of the *GATA1* promoter in K562 H3K27ac HiChIP at 1 kb resolution. CRISPRi-validated regulatory regions in K562 cells are indicated^5^. **(C)** Correlation of *MYC* K562 H3K27ac HiChIP signal with max CRISPRi score within the HiChIP 10 kb window. **(D)** Interaction profiles of the *MYC* promoter in GM12878 and My-La H3K27ac HiChIP at 10 kb resolution. T cell H3K27ac ChIP-seq and ATAC-seq are from Naïve T cells. **(E)** (Top) CRISPRi validation in GM12878 cells of GM12878 and My-La-biased *MYC* enhancers. (Bottom) *MYC* RNA levels by qRT-PCR and cell growth rates in CRISPRi GM12878 cells targeted to cell type-biased enhancers, the *MYC* promoter, and a non-targeting negative control (n = 3). * corresponds to p-value < 0.05, ** to p-value < 0.01, *** to p-value < 0.001, **** to p-value < 0.0001, and n.s. to not significant. The box extends from the 25^th^ to 75^th^ percentiles with a line representing the median, and the whiskers go the minimum and maximum values. **(F)** Interaction profile of the *CD69* promoter in Jurkat H3K27ac HiChIP at 5 kb and 1 kb resolutions. The 1 kb profile is focused on the window of the CRISPRa tiling screen. CRISPRa-validated regulatory regions in Jurkat cells are indicated^26^. **(G)** (Top) CRISPRa validation in Jurkat cells of *CD69* distal enhancers. (Bottom) *CD69* RNA and protein levels in CRISPRa Jurkat cells targeted to distal enhancers, the *CD69* promoter, the *KLRF2* promoter as a locus negative control, and a non-targeting negative control (n = 2).

We found the enhancer landscapes of the *MYC* promoter to be highly cell-type specific. v4C analysis of the *MYC* promoter in GM and My-La (CD4+ T cell leukemia) cells showed dramatically different regulatory interactions with the promoter compared to K562 cells (Fig. 2d). To validate EIS specificity, we performed CRISPRi experiments in GM cells using sgRNAs targeting enhancers identified in either GM or My-La HiChIP maps as well as a positive control sgRNA targeting the *MYC* promoter and a negative control sgRNA targeting lambda phage sequence (Fig. 2e). As expected, we found that CRISPRi of GM, but not My-La, enhancers impacted *MYC* expression and cell growth in GM cells (Fig. 2e).

Finally, we focused on the *CD69* locus, where a high-resolution CRISPRa screen identified three enhancers upstream of the transcription start site^26^. These sites were also identified by Naïve T cell H3K27ac HiChIP. Moreover, HiChIP identified four additional distal enhancers that were outside the region spanned by the sgRNA tiling array (**Fig. 2f and Supplementary Fig. 5**). To functionally validate these novel enhancers, we performed CRISPRa experiments in Jurkat cells with sgRNAs targeting these enhancers, the *CD69* promoter, the *KLRF2* promoter as a locus negative control, and a non-human genome-targeting negative control. We observed a significant increase in *CD69* RNA and protein levels in the four HiChIP enhancers compared to negative controls (**Fig. 2g and Supplementary Fig. 5**). Interestingly, two of the four identified novel enhancers were within promoter regions of distant genes. These findings are in line with previous reports that identified widespread distal gene regulatory functions of promoters genome-wide^27,28^. Altogether, these results suggest that H3K27ac HiChIP EIS identifies functional regulatory elements, and that enhancers that regulate a gene of interest can differ significantly between cell-types.

### Landscape of enhancer interactions in primary T cells

We examined global features of the enhancer connectome associated with cellular differentiation from Naïve T cells to either TH17 cells or Treg cells. We identified a total of 10,706 high confidence loops in the union set of the three cell types (**Supplementary Table 2**). Analysis of loop read support between biological replicates demonstrated high reproducibility (**Supplementary Fig. 4c**), and ~91% of loop anchors were associated with either a promoter or enhancer^29^, as expected, with a median distance of 130 kb (**Supplementary Fig. 6a,b**). Importantly, high-resolution E-P connectivity maps revealed several features that could not be discerned from 1D epigenomic data (i.e. H3K27ac ChIP-seq or ATAC-seq; Fig. 3a). These features included: (i) ‘enhancer skipping’: enhancers that have stronger EIS with a more distal target promoter, (ii) higher order structures such as ‘enhancer cliques’ (related to loop cliques^30^): multiple regulatory elements that have strong EIS with a single target promoter, (iii) promoter to promoter interactions^13,31^, and (iv) ‘enhancer switching’: enhancers that exhibit differential EIS with a target promoter in a cell type-specific manner (Fig. 3a).

**Fig. 3.**
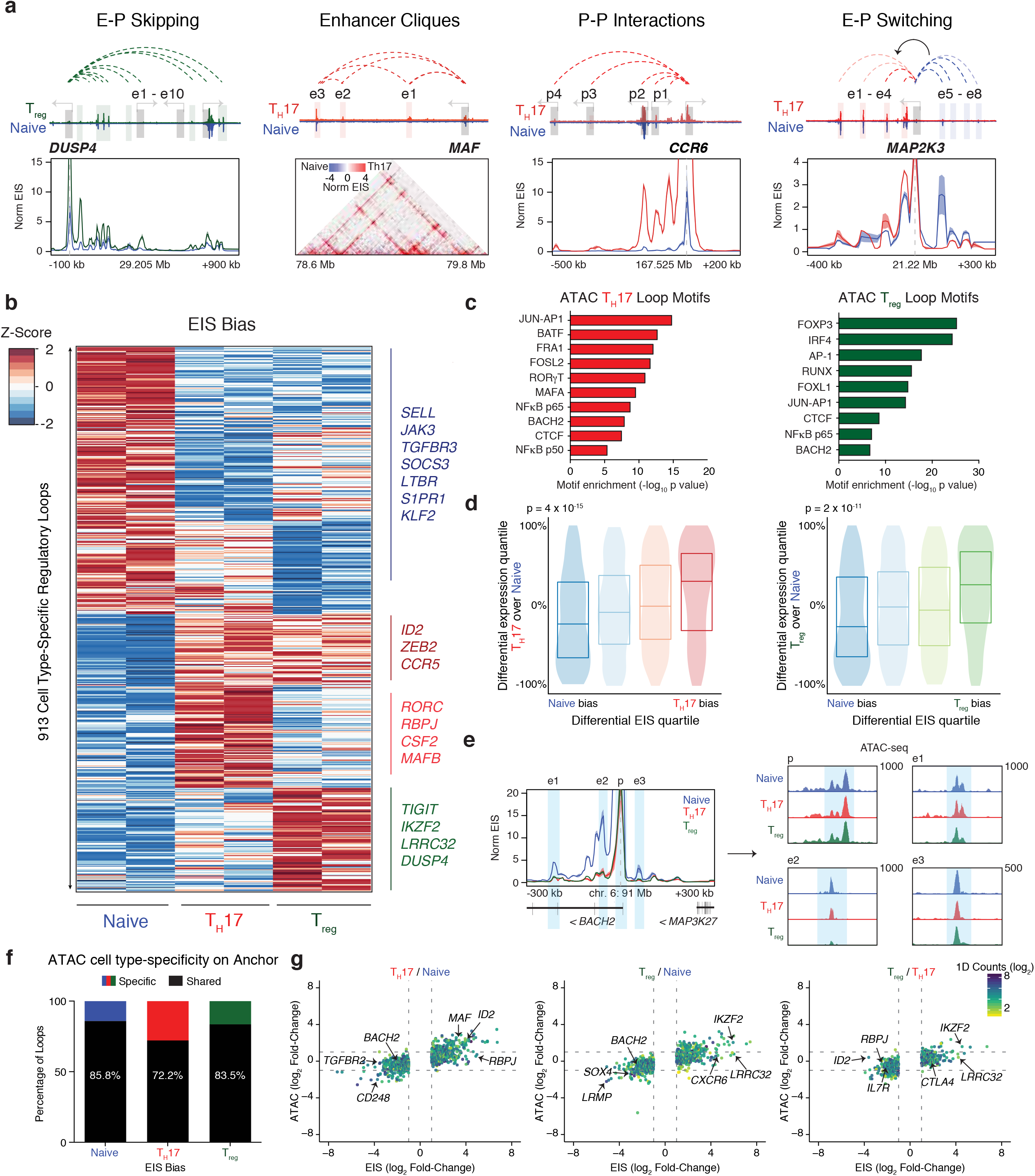
Dynamic 3D enhancer landscapes in T cell differentiation. **(A)** Conformational features observed by H3K27ac HiChIP. **(B)** HiChIP EIS in 913 differential interactions identified in T cell subtypes. Interactions are clustered by cell-type specificity. **(C)** Cell-type specific motif identification from ATAC-seq peaks in biased EIS anchors. **(D)** EIS bias quartiles for Naïve to T_H_17 and Naïve to T_reg_ differentiation, with corresponding differential RNA gene expression rankings. **(E)** Proportion of ATAC-seq peaks within HiChIP differential interaction anchors that are cell-type specific (log_2_ fold change >1) or shared across all three subtypes. **(F)** Interaction profile of the *BACH2* promoter at 5 kb resolution, demonstrating shared accessibility signal at Naïve-biased EIS. **(G)** Global correlation of EIS and ATAC-seq fold-change in different T cell subset pairwise comparisons.

We found that EIS contacts were very cell type-specific. After quantile-quantile normalization of contact reads at high-confidence loops (correcting for false positives caused by 1D fragment visibility; Methods), we focused on the top and bottom 5% of EIS ranked by cell-type bias for each pair-wise comparison (**Supplementary Figs. 6c-g and 7, Supplementary Tables 3-4**). Cell type-specific enhancer loop anchors revealed genes encoding canonical T cell subtype TFs and effector molecules (**Fig. 3b, Supplementary Figs. 8 and 9**). Deeper v4C analysis of shared and cell type-specific loci pinpointed regulatory elements interacting with each gene promoter of interest as well as local conformational landscape changes (**Supplementary Figs. 8 and 9**). TF motifs located within cell type-specific loop anchors were enriched for TFs known to drive T cell subtype differentiation and nominated novel TFs involved in regulation (Fig. 3c). Furthermore, cell type EIS bias was associated with differential expression of genes located within corresponding EIS anchors for the same cell type (Spearman’s rho = 0.242 and 0.207; p-value = 4 x 10^-15^ and 2 x 10^-11^; Fig. 3d).

Cell type-specific EIS may be driven by cell type-specific enhancer activation (based on H3K27ac ChIP-seq) or stable enhancer activation with cell type-specific looping (Hi-C) in a gene specific manner. We first examined H3K27ac ChIP-seq at differential EIS anchors and found that many biased H3K27ac HiChIP interactions also exhibited biased ChIP-seq signal, as expected. 58.5% of Naïve-biased loops contain at least one Naïve-biased ChIP-seq peak (log_2_ fold change > 1) located on the anchors. Similarly, 66.7% of TH17-biased and 67.8% of T_re_g-biased interaction anchors were cell type-specific in 1D (**Supplementary Fig. 10a**). Therefore, while on average ~64% of the differential EIS corresponded to change in 1D data, ~36% were likely also driven by change in 3D chromatin loop strength. To further assess the contribution of cell type-specific 3D signal to EIS, we examined HiChIP 1D signal at differential EIS anchors. We found that HiChIP 1D signal correlated better with ChIP-seq signal than EIS, with a higher likelihood of differential ChIP-seq signal overlapping differential HiChIP 1D signal compared to 3D, suggesting EIS bias is in part driven by 3D changes (**Supplementary Fig. 10b**).

We asked whether the integration of reference cell line Hi-C data with primary T cell H3K27ac ChIP-seq could recapitulate HiChIP EIS in primary T cells. We binned Gm Hi-C loops with increasing primary T cell ChIP-seq signal at loop anchors and then determined the overlap of loops in each bin with loops derived from H3K27ac HiChIP. As expected, increased ChIP-seq signal at the Hi-C anchors led to increased overlap with the HiChIP loops. However, the overlap was lower in all T cell subtypes compared to the same analysis performed using GM HiChIP data. These observations demonstrate that cell-type specific 3D interactions can impact EIS independent of differences in 1D ChIP-seq signal (**Supplementary Fig. 10c**). Similarly, previously generated enhancer-promoter maps obtained from bulk T cells did not identify T cell subtype-specific interactions obtained using H3K27ac HiChIP. To assess the unique information obtained through cell type-specific interaction maps, we compared promoter Capture Hi-C maps in bulk CD4+ T cells to H3K27ac HiChIP maps in Naïve, T_H_17, and Treg cells^14^. Strikingly, the most cell type-specific loops in T_H_17 and Treg (16-fold enriched) demonstrated a low discovery rate in promoter Capture Hi-C T cells (11.83% in 415 loops and 13.83% in 373 loops, respectively; **Supplementary Fig. 10d**). Many of these subset-specific interactions included genomic loci encoding functionally important effector genes, such as *LRRC32*. The *LRRC32* locus contains T_re_g-specific loops that are neither visualized in HiChIP maps from Naïve or T_H_17 cells nor in bulk CD4+ promoter Capture Hi-C maps (**Supplementary Fig. 10e**). Since primary human T_H_17 and T_reg_ cells are present in human blood with low frequency, it would also be challenging to generate subset-specific promoter Capture Hi-C maps with published promoter Capture Hi-C protocols. In summary, EIS is derived from a combination of 1D ChIP-seq and 3D interaction signal and cannot be accurately predicted from 3D maps in reference cell lines or unsorted primary cell datasets.

Cell type-specific EIS can occur at sites of shared chromatin accessibility. Paired chromatin accessibility profiles by Assay of Transposase-Accessible Chromatin by sequencing (ATAC-seq)^32^ from each T cell subset revealed most cell type-specific loop anchors had equivalent chromatin accessibility across all three cell types (Fig. 3e-g). To illustrate this finding, we examined the *BACH2* promoter, which exhibits shared chromatin accessibility at enhancers, but increased EIS in Naïve cells (Fig. 3e). Globally, only 14.2%, 27.8%, and 16.5% of Naïve-, T_H_17-, and T_re_g-biased loops, respectively, contained at least one biased ATAC-seq peak (log_2_ fold change > 1) located on the anchors. Furthermore, the majority of cell type-specific TF motifs were observed in shared ATAC-seq peaks within differential interactions, highlighting that these regions are functioning in T cell differentiation (Fig. 3f-g). Altogether, these results suggest that in highly related – yet functionally distinct – cell types, a portion of transcriptional control is achieved through differential chromosome looping, rather than differential chromatin accessibility. This finding is consistent with previous studies which demonstrated that T cell subset-specific TFs, such as Foxp3, act predominantly at pre-accessible chromatin sites to establish subset-specific gene expression^33^.

### Enhancer interactions link disease variants to target genes

The high specificity of EIS enabled us to identify putative target genes of autoimmune disease risk loci in functionally relevant T cell subsets. To achieve this, we used a previously described list of putatively causal variants associated with 21 autoimmune diseases, known as PICS SNPs, which were fine-mapped based on dense genotyping data^25^. We determined that PICS autoimmune SNPs were significantly enriched in T cell loop anchors, with specific autoimmune diseases showing greater than 5-fold enrichment compared to a shuffled control loop set (**Supplementary Fig. 11**). Next, we constructed a set of all possible connections between autoimmune risk SNPs and TSS within 1 Mb and measured the EIS for each SNP-TSS pair (Fig. 4a). We aggregated these signals to determine the overall interaction activity in each T cell subtype in each disease (Fig. 4b). We observed high interaction strength enrichments and cell type specificity in autoimmune disease SNPs, but low enrichment and cell specificity in non-immune traits (Fig. 4b). To further visualize HiChIP bias in shared or differential enhancers, we analyzed SNP-TSS interactions grouped by their presence near H3K27ac ChIP-seq peaks (**Supplementary Fig. 12a,b**). We observed a large number of active SNP-TSS pairs that were present in regulatory regions that were shared between T effector cell types (T_reg_ and T_H_17), while relatively less EIS signal was observed in SNPs located in cell-type specific enhancers, supporting the concept that many autoimmune disease variants impact common T cell effector/activation pathways^25,34^. Notably, SNPs present in enhancers shared across all three cell types could still be distinguished by HiChIP bias (**Supplementary Fig. 12a,b**). For example, although we could not detect cell type bias at risk loci for Alopecia Areata using H3K27ac ChIP-seq (**Supplementary Fig. 12a,b and ref. 3**), H3K27ac HiChIP identified increased SNP-TSS activity in T_reg_ cells among shared T cell enhancers, consistent with several studies identifying the crucial role of this cell type in disease pathogenesis^35^. Importantly, autoimmune signal enrichments were not readily apparent from 1D H3K27ac ChIP-seq peaks, aggregated ChIP-seq signal within the TAD containing the SNP, nor cell line H3K27ac HiChIP datasets (**Fig. 4b and Supplementary Fig. 12c**). Therefore, examining 3D disease variant interactions may capture cell type biases more robustly than 1D epigenomic data. Finally, to validate our findings with an orthogonal dataset, we performed SNP-TSS EIS analysis on an overlapping set of autoimmune disease-associated SNPs obtained from the NHLBI GRASP catalog and observed similar enrichments of specific T cell subsets (**Supplementary Fig. 12d**).

**Fig. 4.**
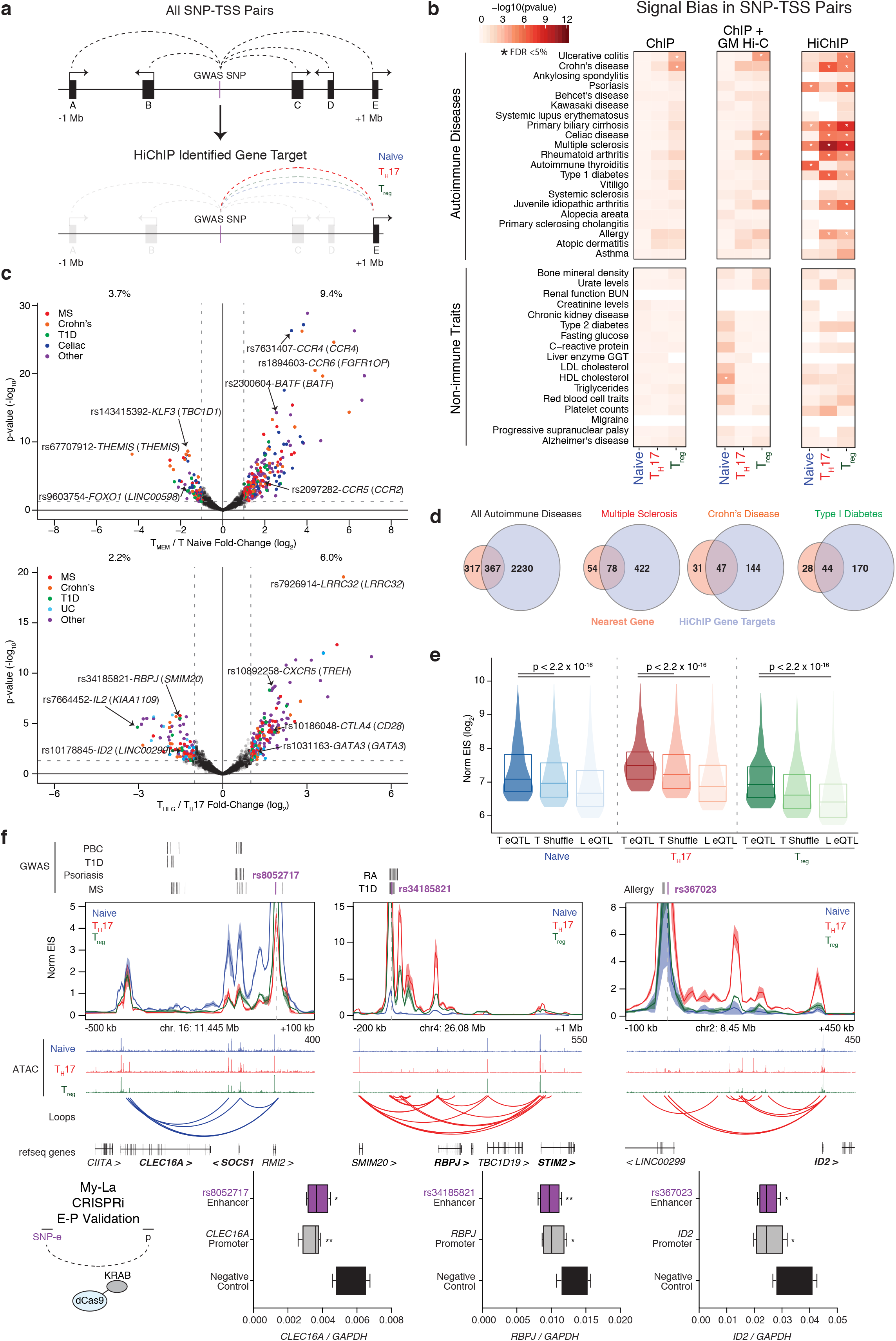
HiChIP identifies cell type-specificity and target genes of autoimmune diseases variants. **(A)** Generation of a loop set between all autoimmune SNPs and gene promoters within a 1 Mb region. **(B)** H3K27ac ChIP and HiChIP signal bias in T cell subtypes for SNP-TSS pairs. For each bin, PICS SNPs are tagged by H3K27ac only in the concordant cell type for the bias tested. SNPs are grossly divided into associations with autoimmune diseases or control, non-immune traits. **(C)** EIS Bias of SNP-TSS loops (with nearest gene annotated) in T_H_17 and Treg subsets versus Naïve, and T_H_17 versus T_reg_. **(D)** Number of HiChIP gene targets versus nearest gene predictions for all looping nongenic autoimmune SNPs as well as SNPs for specific diseases. **(E)** Global validation of HiChIP SNP gene targets. Synthetic SNP-TSS pairs were generated from each CD4+ eQTL SNP to its associated gene and compared to both a distance-matched shuffled SNP-TSS pair and a liver eQTL SNP-TSS pair. **(F)** HiChIP target gene RNA levels by qRT-PCR in CRISPRi My-La cells targeted to SNP-containing enhancers of interest, as well as positive control sgRNAs to the HiChIP target promoters and a non-targeting negative control (n = 3). * corresponds to p-value < 0.05, ** to p-value < 0.01, *** to p-value < 0.001, **** to p-value < 0.0001, and n.s. to not significant. The box extends from the 25^th^ to 75^th^ percentiles with a line representing the median, and the whiskers go the minimum and maximum values.

We leveraged HiChIP to identify potential gene targets of intergenic SNPs, which have classically been paired to the nearest neighboring gene. We overlapped the SNP-TSS pairs with loops to call a discrete set of target pairs. We then performed differential analysis on the SNP-TSS loops to ascertain bias for specific T cell subsets (**Fig. 4c and Supplementary Table 5**). Examples of biased SNP-TSS pairs included *FOXO1* in Naïve T cells (rs9603754), *BATF* (rs2300604) in Memory T cells, *CTLA4* (rs10186048) in Treg cells, and *IL2* (rs7664452) in T_H_17 cells (**Fig. 4c and Supplementary Table 5**). Next, we sought to characterize the connectivity landscape of the SNP-TSS loops. We identified an average of 1.75 gene targets per autoimmune SNP (ranging from 0 to over 10 target genes), while non-immune traits did not demonstrate an increase in targets (0.33 genes per SNP; **Supplementary Fig. 12e**). For 684 autoimmune intergenic SNPs, we identified a total of 2,597 HiChIP target genes, representing a four-fold increase in target genes for known disease SNPs (Fig. 4d). Only 367 (~14%) of all targets were the nearest gene to the SNP, while approximately ~86% of SNPs skipped at least one gene to reach a predicted target TSS (**Supplementary Fig. 12e**). Furthermore, approximately ~45% of SNP to HiChIP target interactions had increased signal compared to the same SNP to nearest gene, despite distance biases.

### Target gene validation by eQTL and CRISPRi

HiChIP enhancer-target gene interactions can be validated using previously identified point mutations that alter expression at distantly located genes in T cells—i.e. expression quantitative trait loci (eQTL)^36^. For example, the celiac disease-associated SNP rs2058660 impacts the expression of the inflammatory cytokine receptor genes *IL18RAP, IL18R1, IL1RL1*, and *IL1RL2*, which are known regulators of intestinal T cell differentiation and response^37^. HiChIP EIS revealed contacts between rs2058660 and each of these predicted gene promoters (**Supplementary Fig. 13a**). Similarly, the Crohn’s disease risk variant rs6890268 and the multiple sclerosis (MS) risk variant rs12946510 impact the expression of *PTGER4* and *IKZF3*, respectively, and H3K27ac HiChIP also demonstrated clear contacts between these SNPs and their predicted promoter (**Supplementary Fig. 13a**). Globally, HiChIP contact signal was increased in eQTLs in T cells compared to a distance-matched background loop set (p-value < 2.2 x 10^-16^; Fig. 4e) or to eQTLs identified in an unrelated cell type (liver; p-value < 2.2 x 10^16^). The overlap of HiChIP and eQTL loci provides support for chromosome interactions as a physical basis for distal eQTLs^10-12^ and further validates the HiChIP approach to assign enhancer-target gene relationships.

We next sought to directly validate HiChIP SNP-gene targets using CRISPRi in My-La cells. First, we focused on three loci of interest in primary T cells and then confirmed that the SNP-TSS loops were also present in My-La cells (**Fig. 4f and Supplementary Fig. 13b**). We then targeted sgRNAs to these SNP-containing enhancers, as well as positive control sgRNAs to the HiChIP target gene promoters and a negative control sgRNA targeting lambda phage sequence. As expected, we observed a significant reduction of RNA levels in the HiChIP target genes upon CRISPRi of its SNP-containing enhancer (Fig. 4f).

### Fine-mapping of disease-associated DNA variants

Since SNP-TSS HiChIP signal is capable of identifying target genes of candidate SNPs, we asked whether TSS-SNP HiChIP signals could also be used to nominate functional causal variants within haplotype blocks in a reciprocal manner. We first performed a proof-of-principle analysis using fine-mapped SNPs associated with inflammatory bowel disease (IBD)^38^ or Type 1 Diabetes (T1D)^39^ as well as high confidence PICS SNPs and examined EIS from putatively causal SNPs to all gene promoters within 300 kb. EIS from putatively causal SNPs to gene promoters was significantly higher than EIS from a distance-matched set of SNPs within the same LD block to gene promoters (p-value = 2.4 x 10^-15^, 8.7 x 10^-8^, 3.9 x 10^-3^ for IBD fine-mapped SNPs, T1D fine-mapped SNPs, and high confidence PICS, respectively; **Fig. 5a and Supplementary Fig. 14a**). Next, we assessed the fine-mapping ability of HiChIP EIS at individual loci of interest. We focused on IBD-and MS-associated SNPs neighboring the *PTGER4* and *SATB1* loci and performed v4C analysis anchored at the gene promoters. We calculated EIS signal at 1 kb resolution and identified specific regions within the linkage disequilibrium (LD) blocks that contained the highest EIS to the target promoters, positioning the likely causal SNPs within these regions (**Fig. 5b and Supplementary Fig. 14b**). For example, at the *PTGER4* locus (Fig. 5b), the ~160 kb genomic interval spanned by LD SNPs in association with Crohn’s disease is refined to two bins of 3kb and 4kb, which both contain PICS SNPs.

**Fig. 5.**
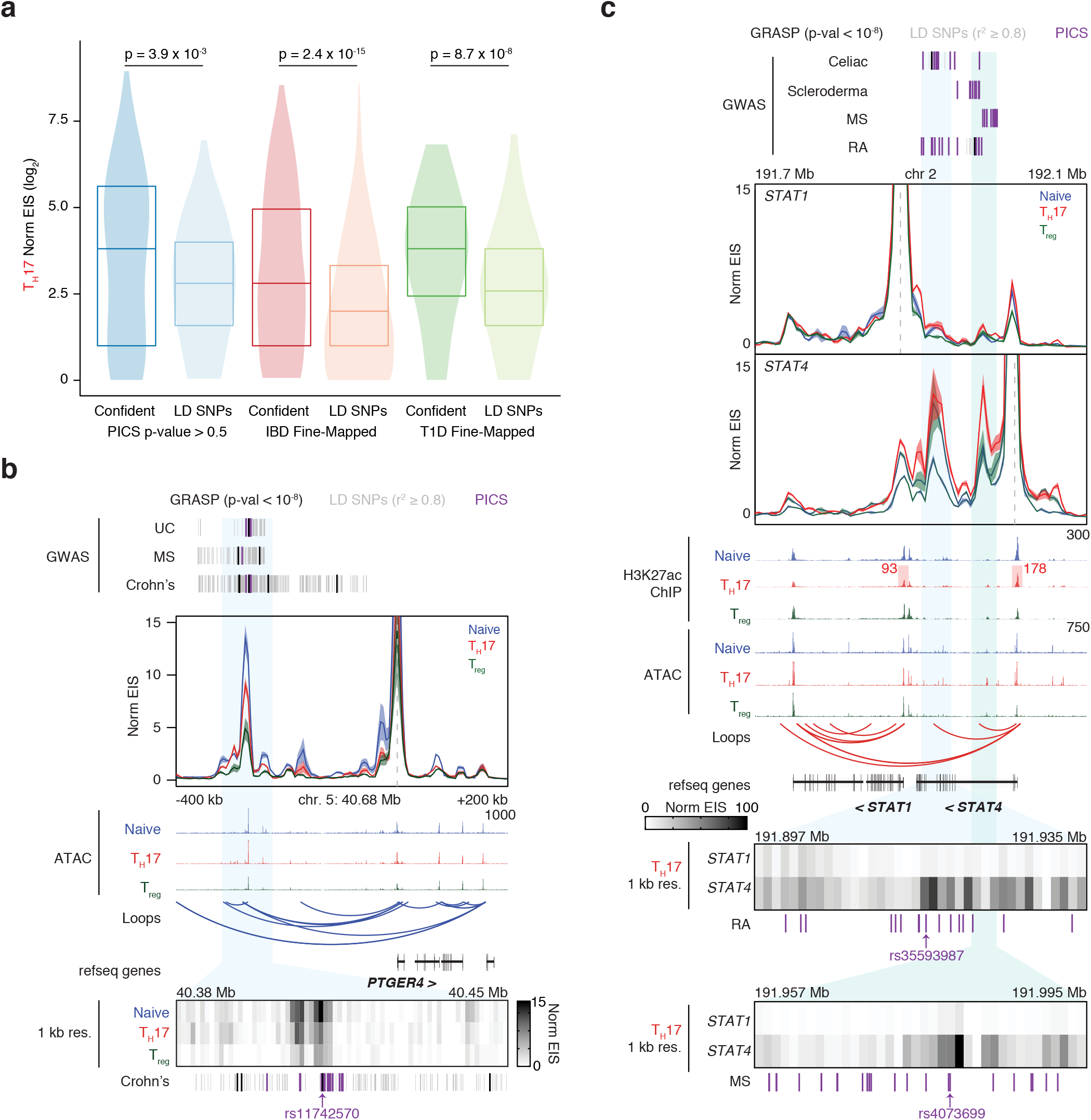
Fine-mapping of GWAS variants using H3K27ac HiChIP. **(A)** Global validation of HiChIP signal at putatively causal SNPs versus corresponding SNPs in LD (r^2^ > 0.8) for T_H_17 cells. SNP-TSS pairs were generated from published fine-mapped datasets, compared to a distance-matched SNP-TSS pair set in the same LD block. **(B)** Interaction profile of the *PTGER4* promoter, and a 1 kb resolution visualization of the SNP-containing enhancer of interest. LD SNPs (r^2^ ≥ 0.8) correspond to GRASP SNPs (genome-wide significance p-value < 10^-8^). The highlighted SNP was identified in both the high confidence PICS and GRASP datasets. **(C)** Interaction profiles of the *STAT1* and *STAT4* promoters, with 1 kb resolution visualizations of the SNP-containing enhancers of interest. Highlighted are 1D signal contributions at the *STAT1* and *STAT4* promoters. Highlighted SNPs are PICS closest to focal EIS to *STAT4*.

We asked whether complex disease-associated loci containing more than one gene could be fine-mapped using HiChIP. We focused on two disease-associated enhancers in between the *STAT1* and *STAT4* gene promoters (Fig. 5c). These two genes encode transcription factors with distinct roles in immune regulation. Signal transducer and activator of transcription 1 (STAT1) is critical for type I IFN and IFNγ signaling, whereas STAT4 induces T_H_1 differentiation and IFNγ expression^40^. We investigated bias of these enhancers to *STAT1* and *STAT4* and found that, despite comparable linear distance and 1D signal at the promoters, the enhancers were significantly biased to interact with *STAT4*. Next, we fine-mapped the disease associated SNPs within this locus using 1 kb resolution EIS from the *STAT4* promoter, and narrowed down candidate functional variants within the two enhancers (Fig. 5c). In summary, HiChIP EIS can nominate functional causal variants within haplotype blocks, and two-way analysis of target gene identification from an enhancer of interest and high-resolution interaction maps of that enhancer with its target gene can be used to fine-map disease-associated loci containing several candidate genes.

### Allelic target gene bias of cardiovascular disease variants

Finally, we asked whether this approach could be applied broadly to other categories of human disease, and whether we could directly test SNP-TSS associations using allele-specific HiChIP. We generated high-resolution E-P maps from primary human coronary artery smooth muscle cells (HCASMC), which can be used to inform variants linked to cardiovascular diseases^41^. First, to validate cell type specificity, we examined the *TCF21* gene promoter, a transcription factor required for the differentiation of HCASMC^42^ and observed enrichment in HCASMC EIS relative to Naïve T cells (Fig. 6a). We next examined the 9p21.3 locus, which harbors risk associations with several cardiovascular disorders^43-45^. We found that the promoters of all three genes in the locus interact with one another and with CAD variant-containing enhancers located approximately 100 kb upstream of the *CDKN2B* promoter (**Supplementary Fig. 15**). We then generated SNP-TSS target lists using CAD SNPs identified in the CARDIoGRAMplusC4D study^46^. We again performed differential analysis on the SNP-TSS loops to ascertain bias for HCASMC versus Naïve T cells (Fig. 6b). Overall, 75.1% of biased HCASMC SNP-TSS pairs were CAD SNPs, while only 5.5% of Naïve T cell biased SNP-TSS pairs were CAD SNP-TSS loops. Next, we examined the connectivity of the HCASMC SNP-TSS contacts and identified 1,062 gene targets, of which only 120 (~11%) mapped to the nearest gene. Furthermore, approximately 89% skipped at least one gene to reach a predicted target TSS, and 64% of SNPs were mapped to more than a single gene target.

**Fig. 6.**
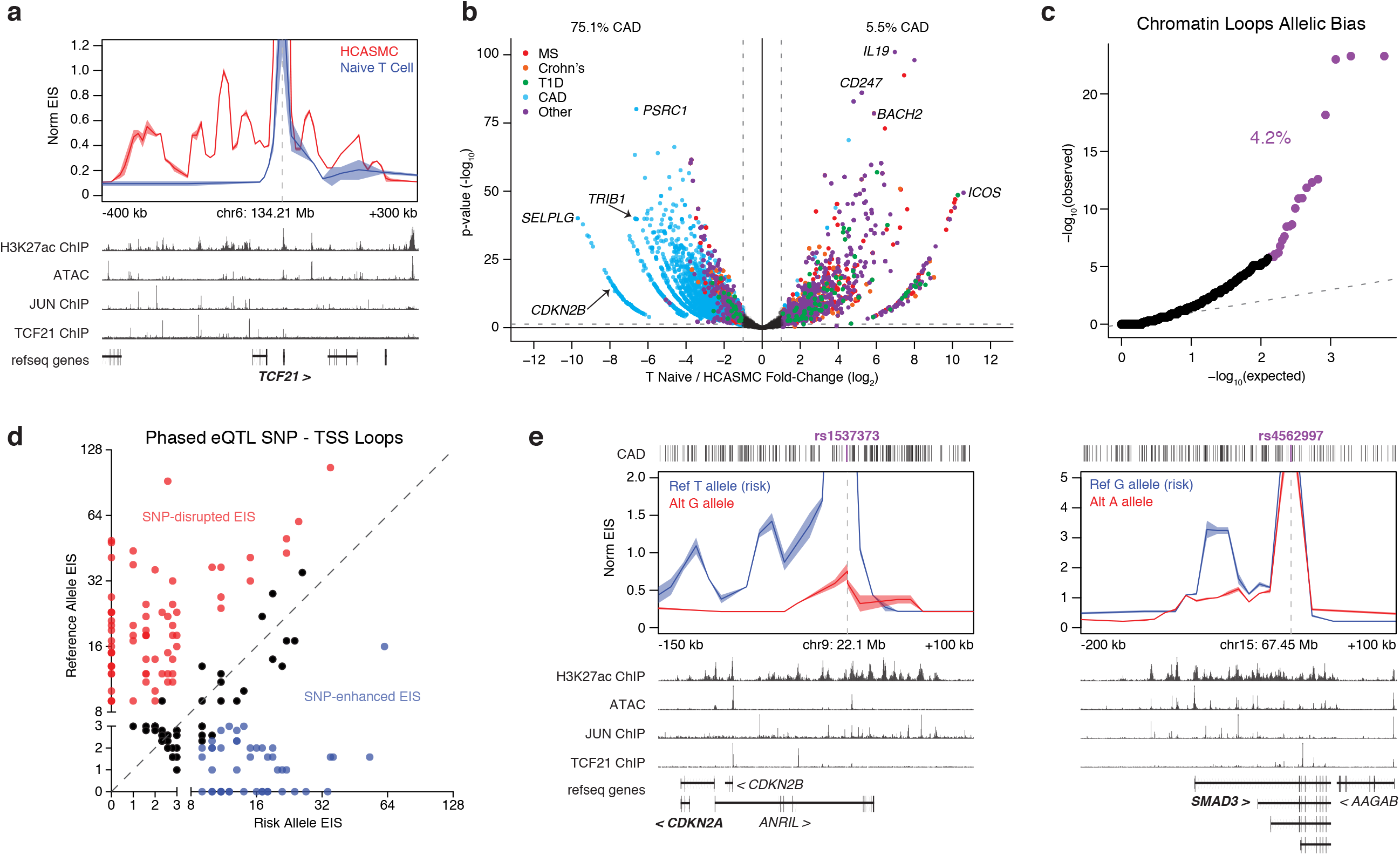
HiChIP identifies allelic bias to target genes for cardiovascular disease risk variants. **(A)** Interaction profile of the *TCF21* gene promoter for H3K27ac HiChIP of HCASMC and Naïve T cells. **(B)** EIS bias between HCASMC and Naïve T cells in a union set of CARDIoGRAMplusC4D CAD and PICS autoimmune SNP-TSS loops. **(C)** Q-Q plot of allelic EIS imbalance in high confidence loops. Allelic mapping biased loops were identified through simulation and removed prior to EIS analysis. **(D)** EIS bias between CAD risk variants and their alternative alleles to eQTL associated target genes. **(E)** Allele-specific HiChIP interaction profiles at the 9p21.3 and *SMAD3* loci at 10 kb resolution in order to examine the functional consequence of a risk variant compared to its alternative allele.

We took advantage of genome phasing information in HCASMC to measure E-P interactions at allele-specific CAD SNPs, allowing us to examine the functional consequence of a risk variant compared to its alternative allele in the same nucleus. First, 4.2% of high confidence loops in HCASMC with no observed mapping bias in the anchors exhibited significant allelic bias (FDR < 0.05, Fig. 6c), consistent with frequency of allelic imbalance of RNA expression and prior evidence of allele-specific regulation of specific E-P interactions^47,48^. We leveraged this global E-P allelic bias to examine the effect of a risk variant compared to its control alternative allele for a set of CAD-associated SNP-target gene pairs (Fig. 6d)^49^. We found that many risk alleles disrupt enhancer-target gene interactions, but a subset of pathogenic SNPs increased enhancer-target gene interaction. At CAD risk variant rs1537373 in the 9p21.3 locus, the risk allele (T) showed increased EIS to the *CDKN2A* promoter as well as an additional enhancer within the lncRNA *ANRIL* relative to the reference allele (G) (Fig. 6e). We further observed increased EIS of the CAD risk variant rs4562997 to an additional *SMAD3* enhancer 10 kb downstream of the TSS (Fig. 6e). The ability to resolve enhancer connectomes of the risk and reference alleles in the same nucleus demonstrates that the mutated base in the risk allele suffices to alter enhancer looping in *cis* in disease-relevant primary cells.

## DISCUSSION

Here, we developed an approach to define the high-resolution landscape of E-P regulation in primary human cells. We find that E-P contacts are highly dynamic in related cell types and often involve genomic elements with shared accessibility. Accordingly, many complex features of the 3D enhancer connectome cannot simply be predicted from 1D, which demonstrates that mapping conformation in primary cells can identify novel regulatory connections underlying gene function in human disease. We take advantage of this principle to chart the connectivity of autoimmune and cardiovascular GWAS SNPs and link SNPs to hundreds of potential target genes. Although non-genic SNPs have previously been paired with their closest neighboring gene, we find that the majority of these variants can engage in long-distance interactions, including skipping several promoters to predicted target genes, connecting to multiple genes, or acting in concert with enhancer cliques to contact a single gene. Further use of this approach will help to clarify hidden mechanisms of human disease that are driven by genetic perturbations in non protein-coding DNA elements, which can now be linked to their cognate gene targets in primary cells.

## ACKNOWLEDGEMENTS

We thank members of the Chang and Greenleaf laboratories for helpful discussions and Justin Tumey for artwork. This work was supported by National Institutes of Health (NIH) P50HG007735 (H.Y.C. and W.J.G.) and U19AI057266 (W.J.G), Human Frontier Science Program (W.J.G.), Rita Allen Foundation (W.J.G.), and the Scleroderma Research Foundation (H.Y.C). M.R.M. and E.A.B. acknowledge support from the National Science Foundation Graduate Research Fellowship. A.T.S. is a Cancer Research Institute Irvington Fellow supported by the Cancer Research Institute. B.G.G. was supported by an Ō-AstraZeneca Postdoctoral Fellowship. M.R.C. is supported by a grant from The Leukemia & Lymphoma Society Career Development Program. J.E.C. was supported by the Li Ka Shing Foundation and the Heritage Medical Research Institute. W.J.G. and A.M. are Chan Zuckerberg Biohub investigators. A.M. serves as an advisor to Juno Therapeutics and PACT Therapeutics and the Marson lab has received sponsored research support from Juno Therapeutics and Epinomics. A.M. and J.E.C. are founders of Spotlight Therapeutics. Sequencing was performed by the Stanford Functional Genomics Facility (NIH S10OD018220). We thank Agilent Technologies for generating oligo pools for cloning of the CRISPRa gRNAs. We thank the UC Berkeley High Throughput Screening Facility and Flow Cytometry Facility. H.Y.C. and W.J.G. are founders of Epinomics and members of its scientific advisory board.

## AUTHOR CONTRIBUTIONS

M.R.M, A.T.S., W.J.G., and H.Y.C. conceived the project. M.R.M., A.T.S., J.T., and R.L. performed all genomics assays with help from T.N., M.R.C., N.S., and R.A.F. A.T.S. performed all sorting for experiments. B.G.G., S.W.C., M.R.M., M.L.N., K.R.K., and D.R.S., performed all CRISPR validation experiments. E.A.B., C.D., M.R.M., and J.X. analyzed HiChIP data. J.G., A.T.S., and Y.W. analyzed ATAC-seq data. A.J.R. and P.G.G. analyzed GWAS SNPs in HiChIP data. A.K., P.A.K., A.M., J.E.C., T.Q., W.J.G., and H.Y.C guided experiments and data analysis. M.R.M, A.T.S, E.A.B, C.D., W.J.G, and H.Y.C. wrote the manuscript with input from all authors.

## METHODS

### Human Subjects

This study was approved by the Stanford University Administrative Panels on Human Subjects in Medical Research, and written informed consent was obtained from all participants.

### Cell Culture and Primary T cell Isolation

Mouse ES cells (v6.5, Novus Biologicals: NBP1-41162) were cultured in Knockout DMEM (Gibco) + 15% FBS and leukemia inhibitory factor (LIF, Millipore) to 80% confluence. GM12878 (Coriell), Jurkat, and My-La (CD4+) cells (ATCC) were grown in RPMI 1640 (Gibco) with 15% FBS to a concentration of 500,000 to 1 million cells per mL. Normal donor human peripheral blood cells were obtained fresh from AllCells. CD4+ T cells were enriched from peripheral blood using the RosetteSep Human CD4^+^ T Cell Enrichment Cocktail (StemCell Technology). For CD4^+^ T helper cell subtypes, Naïve T cells were sorted as CD4^+^CD25^-^CD45RA^+^, TH17 cells were sorted as CD4^+^CD25^-^ CD45RA^-^CCR6+CXCR5^-^, and T_reg_ cells were sorted as CD4+CD25+CD127^lo^ Antibodies used for FACS included: PerCP/Cy5.5 anti-CD45RA (Biolegend 304122), Brilliant Violet 510 anti-CD127 (Biolegend 351331), APC/Cy7 anti-CD4 (Biolegend 344616), PE anti-CCR6 (Biolegend 353410), FITC anti-CD25 (Biolegend 302603), Brilliant Violet 421 anti-CXCR3 (Biolegend 353715), and BB515 anti-CXCR5 (BD Biosciences 564625). For HiChIP experiments, 500,000 - 1 million cells were sorted into RPMI + 10% FCS. For ATAC-seq experiments, 55,000 cells were sorted into RPMI + 10% FCS. Post-sort purities of > 95% were confirmed by flow cytometry for each sample.

Primary human coronary artery smooth muscle cell (HCASMC) line derived from a normal human donor heart was purchased from Cell Applications, Inc. (350-05A) and cultured in smooth muscle growth medium (Lonza, CC-3182) supplemented with hEGF, insulin, hFGF-b, and 5% FBS. Cells were grown according to Lonza's instructions.

### Cell Fixation

Detached cell lines or sorted CD4^+^ T cells were pelleted and resuspended in fresh 1% formaldehyde (Thermo Fisher) at a volume of 1 mL formaldehyde for 1 million cells. Cells were incubated at room temperature for 10 min with rotation. Glycine was added at a final concentration of 125mM to quench the formaldehyde, and cells were incubated at room temperature for 5 min with rotation. Finally, cells were pelleted and washed with PBS, pelleted again, and stored at −80 °C or immediately taken into the HiChIP protocol.

### HiChIP

The HiChIP protocol was performed as previously described, using either H3K27ac antibody (Abcam ab4729) or CTCF (Abcam ab70303)^16^ with the following modifications. For primary T cells, we performed HiChIP on as many cells as we could obtain from a blood donation - approximately 500,000 - 1 million cells per T cell subtype per replicate.

We performed two minutes of sonication, no Protein A bead preclearing, used 4 μg of H3K27ac antibody (Abcam ab4729), and captured the chromatin-antibody complex with 34 μL of Protein A beads (Thermo Fisher). Qubit quantification post ChIP ranged from 5 − 25 ng depending on the cell type and amount of starting material. The amount of Tn5 used and PCR cycles performed were based on the post ChIP Qubit amounts, as previously described^16^.

25m cell line libraries were generated as previously described^16^. For low cell number mouse embryonic stem cell samples, we performed two minutes of sonication and no Protein A bead preclearing. Either 4 μg or 2μg of H3K27ac antibody (Abcam ab4729) was used for ChIP in 500k or 100k/50k cells, respectively, and the chromatin-antibody complex was captured with 34 (500k cells) or 20 μL (100k/50k cells) of Protein A beads. Post-ChIP Qubit quantification for the 25m cell samples was approximately 1.5 μg. For lower cell numbers, quantification was 30, 10, and 5 ng for 500k, 100k, and 50k cells, respectively. The amount of Tn5 used and PCR cycles performed were based on the post ChIP Qubit amounts, as previously described.

HiChIP samples were size selected by PAGE purification (300-700 bp) for effective paired-end tag mapping, and therefore were removed of all primer contamination which would contribute to recently reported "index switching" on the Illumina HiSeq 4000 sequencer^50^. All libraries were sequenced on the Illumina HiSeq 4000 to an average depth of 500-600M total reads.

### HiChIP Data Processing

HiChIP paired-end reads were aligned to hg19 or mm9 genomes using the HiC-Pro pipeline^51^. Default settings were used to remove duplicate reads, assign reads to MboI restriction fragments, filter for valid interactions, and generate binned interaction matrices. HiC-Pro filtered reads were then processed into a .hic file using the hicpro2juicebox function. The Juicer pipeline HiCCUPS tool was used to identify high confidence loops^4^ using the same parameters as for the GM12878 *in situ* Hi-C map: hiccups -m 500 -r 5000,10000 -f 0.1,0.1 -p 4,2 -i 7,5 -d 20000,20000 .hic_input HiCCUPS_output. For T cell Juicer loops, performing the default Juicer calls resulted in a high rate of false positives upon visual inspection of the interaction matrix. We therefore called loops with the same HiCCUPS parameters in two biological replicates for each T cell subtype and then filtered loops for those that were reproducibly called in both replicates. In addition, we removed all loops greater than 1 Mb.

1D signal enrichment and peak calling were generated from the HiC-Pro filtered contacts file. Intrachromosomal contacts were filtered and both anchors were extended by 75 base pairs. The combined bed file containing both anchors was then used to generate bigwigs for visualization in the WashU Epigenome Browser or call peaks using MACS2.

Allele-specific HiChIP data processing was achieved using HiC-Pro’s allele-specific analysis features^51^. First, HCASMC phasing data^41^ was used to mask the hg19 genome and make indexes. HiC-Pro settings were similar as to described above, with the exception that reads were aligned to the masked genome, and then assigned to a specific allele based on phasing data.

### Interaction Matrices and Virtual 4C Visualization

HiChIP interaction maps were generated with Juicebox using Knight-Ruiz (KR) matrix balancing and visualized using Juicebox software at 500 kb, 25 kb, 10 kb, and 5 kb resolutions as indicated in each analysis^4^. For 1 kb profiles, raw matrix counts were visualized in Java TreeView.

Virtual 4C plots were generated from dumped matrices generated with Juicebox. The Juicebox tools dump command was used to extract the chromosome of interest from the .hic file. The interaction profile of a specific 5 kb or 10 kb bin containing the anchor was then plotted in R. Replicate reproducibility was visualized with the mean profile shown as a line and the shading surrounding the mean representing the standard deviation between replicates. For the HCASMC data we observed low read coverage for allele-specific v4Cs at loci of interest. This is due to the density of SNPs for this genotype, and a low number of reads containing a phased SNP. We therefore could not observe interaction profiles when visualizing separate replicates with the standard deviation. We therefore utilized pseudoreplicates for the HCASMC v4C visualizations^52^.

High confidence Juicer loop calls were loaded into the WashU Epigenome Browser along with corresponding ATAC-seq profiles and publically available H3K27ac ChIP-seq data from the Roadmap Epigenome Project. Browser shots from WashU track sessions were then included in virtual 4C and interaction map anecdotes.

### Reproducibility Scatter Plots and Correlations

Biological and technical reproducibility comparisons between HiChIP experiments were generated by counting reads supporting a set of Juicer loop calls. For biological replicate comparisons, the loop set called from the merged replicates was used. For the comparison in mES cells between low cell number and 25 million cells, the union loop set between the two maps was used. The Pearson correlation between replicates or experiments was calculated from depth-normalized reads using the cor() function in R. Scatter plots were plotted using log-transformed raw reads supporting each loop.

### Promoter/Enhancer Annotations of HiChIP Loops

Promoters were defined as 1 kb regions centered at the TSS, and enhancers were identified as chromHMM enhancers not overlapping with promoters in any cell type. We annotated loop anchors as ‘other’ if the anchor did not contain a promoter or enhancer as defined above.

### Differential Analysis of HiCCUPS Loop Calls

Juicer loop calls from the three T cell subtypes were initially combined into a union set of T cell loops. Loop signal was then obtained for the biological replicates of each T cell subtype. Vanilla coverage square root (VCsqrt) normalized signal for the interaction matrix of each biological replicate using the Juicebox tools dump command. Normalized signal was then assigned to the union loop set in each replicate.

VCsqrt signal per sample was quantile-quantile normalized under the assumption that overall signal was identically distributed across all samples. Following normalization, samples for Naïve, T_H_17 and Treg cells shared Pearson correlations of 0.938, 0.942 and 0.934, respectively. PCA was performed using the prcomp function in R, which demonstrated that the first PC, which exhibited near-identical loadings across the six samples, explained 93% of the variance across the six samples. This was taken to represent the shared signal across cell types. PCs 2-4 explained 2.2%, 2.0% and 1.4%, respectively.

To study cell type-specific looping, the residual signal per loop was taken after projecting the loop onto the unit vector along the diagonal (equal signal per cell type). Cell type-specific and differential looping analysis were performed with the top and bottom 5% of the distributions of either residual signal or differences between cell type residual signals. Hierarchical clustering was performed using the union of all differential loops in these extremes and using 1 minus the Pearson correlation as the distance metric. QQ plots were generated by permuting residuals from the same cell type or individual and summing them together and using this distribution to calculate p values for the observed sums.

In parallel, differential loops were called using edgeR for both the mES and T cell datasets. Again, biological replicate loop signal was obtained across a union set of Juicer loops. We then used edgeR to identify loops with significant changes in signal among pair-wise comparisons (FDR < 0.1, log_2_FC > 1). Importantly, inspection of differential loops identified from the two methods revealed high concordance.

Gene density was calculated from Ensembl gene annotations. GC content was calculated per 10 kb bin using the bedtools nuc function and aggregated as needed. Notably, Spearman correlation between gene density of an entire chromosome and the number of differential loop anchors (rho = 0.914) was much higher than the correlations between the variance in cell type signal per loop anchor and number of genes per 10 kb window (rho = 0.322) and between differential loop anchors and gene density per 100 kb section (rho = 0.083). Correlations between GC content and number of differential loops were similar at both the chromosome (rho = 0.729) and 100 kb bin (rho = 0.148) levels, but while local GC content is likely to confound relative abundance, it is unclear how chromosome-wide GC content could have the same effect.

For mES analysis of H3K27ac- and cohesin-mediated HiChIP, we performed edgeR to obtain the biased loops for each factor. To determine functional bias of the top loops, overlap was determined between edgeR differential loop anchors and relevant ChIP-seq peaks. Smc1a ChIP-seq peaks were obtained from a published dataset^53^. CTCF, RNA Polymerase II, and H3K27ac ChIP-seq peaks were obtained from the mouse ENCODE repository^54^.

### RNA Expression Analysis

Previously generated RNA-seq data^55^ from Naïve, T_H_17 and Treg cells was downloaded as fastqs from ArrayExpress. Illumina adaptors were trimmed using CutAdapt and Ensembl’s cDNA transcripts were quantified using kallisto. Sleuth was used to identify transcripts that were differentially expressed across cell types with FDR controlled at 5%. The mean TPM was calculated per cell type, and TSS differential looping quantiles at genes with with nonzero expression were correlated with differential expression quantiles of the same genes. Only 10 kb segments of the genome that contained a single annotated gene were considered to avoid errors in attribution of looping signal per 10 kb bin. For genes with multiple annotated TSSs the 10 kb bin corresponding to the median TSS was used. Significance was assessed by the cor.test function in R.

### Comparing HiChIP to Hi-C, ATAC-seq, ChIP-seq, and Capture-C datasets

T cell subset ChIP-seq data was obtained from the WashU Roadmap repository. A union set of peaks were called using MACS2 and peaks were quantified using Bedtools intersect. Normalization for ChIP was performed using quantile normalization using “preprocessCore” package in R.

Differential EIS was determined using TMM normalization in the “edgeR” package in R. The significantly differential EIS (log_2_ fold-change > 1 and FDR < 10%) were determined for each pairwise comparison. For each differential EIS, the maximum ATAC/ChIP signal peak was assessed in each 10 kb anchor (to bias against low signal peaks) and then the maximum log_2_ fold-change was compared to the differential EIS.

Differential 1D HiChIP was determined using TMM normalization in the “edgeR” package in R. The significantly differential 1D HiChIP in EIS 10 kb anchors (log_2_ fold-change > 1 and FDR < 10%) were determined for each pairwise comparison. For each differential 1D HiChIP anchor, the maximum ATAC/ChIP signal peak was assessed (to bias against low signal peaks) and then the maximum log_2_ fold-change was compared to the differential 1D HiChIP.

To determine if reference cell line Hi-C data with primary T cell H3K27ac ChIP-seq could recapitulate EIS in primary T cells, GM Hi-C loop anchors were binned with increasing T cell subset ChIP signal as well as GM H3K27ac ChIP-seq signal (Encode). Loop overlap was then determined for the different H3K27ac ChIP signal bins with HiChIP loops.

To compare directly with CD4+ Capture-C data, CHiCAGO loops were called in Naïve, T_H_17, and Treg datasets. CHiCAGO calls were then combined into a union set, and loop signal was obtained for the biological replicates of each T cell subtype using the Juicebox tools dump command. We then used edgeR to identify loops with significant changes in signal among pair-wise comparisons (FDR < 0.05). Total and differential loops were then overlapped with CD4+ Capture-C CHiCAGO data.

### Calculation of Disease-specific GWAS SNP Enrichment in Loop Anchors

We categorized GWAS SNPs into sets relevant to diseases of the immune system and, separately, diseases with no known immune component^25^. For each disease in the immune or non-immune set, we determined the proportion of all GWAS SNPs associated with that disease which overlap the positions of loop anchors based on a union set of loops identified in Naïve, T_H_17, and Treg cells. The ratio of the proportion of immune and non-immune overlaps relative to a shuffled control was reported as the enrichment of immune GWAS SNPs.

### Distance-matched eQTL SNP-TSS Comparisons

We obtained three groups of eQTL SNP-TSS pairs within 1 Mb distance for HiChIP EIS comparisons. The treatment group contains CD4 eQTL-TSS targets. We have two distance-matched groups as control. The first control group contains the CD4 eQTL SNP-random TSS pairs such that the distance between eQTL SNP and random TSS differs by at most 5 kb with the treatment group. The second control group contains liver eQTL SNP-TSS targets that are also distance matched with the treatment group. The random eQTL SNP-TSS pairs were generated by individual chromosome, such that number of control pairs and treatment pairs are the same for every chromosome. In total, there were 158,482 distance-matched eQTL-TSS pairs. We compared the 5 kb resolution EIS among the three eQTL SNP-TSS groups for all three T cell subtypes. Results show that in all the cases, the EIS between CD4 eQTL-TSS targets were significantly higher than the two control groups (p-value < 10^-16^).

### Distance-matched Fine-mapped SNP-TSS Comparisons

We obtained a list of putatively causal SNPs from the PICS SNP list^25^ (PICS probability > 0.5), as well as fine-mapped SNPs associated with inflammatory bowel disease (IBD)^38^ or Type 1 Diabetes (T1D)^39^. Next, we obtained all SNPs in LD with each putatively causal SNP using European linkage disequilibrium blocks determined by all SNPs with an r^2^ ≥ 0.8 with the SNP being considered. For the fine-mapped (T1D/IBD) sets, using SNPs in LD with highly significant GWAS SNPs may mean that there are several SNPs of equal or greater significance in the control set, but we still expect an enrichment relative to the LD block.

We collected all the synthetic pairs between the putatively causal immune-disease related SNPs (IBD, T1D and PICS) and nearby genes within 300 kb distance. To perform the distance-matched EIS comparisons, for each fine-mapped SNP category, we selected the SNP-TSS control pairs which satisfy two constraints: (1) the selected control SNP is positioned at least 5 kb away from the fine-mapped SNP in the same linkage disequilibrium block; (2) the distance of SNP-TSS control pair differs with fine-mapped SNP and the target gene at most 5 kb.

### SNP-TSS Loop Analyses

We obtained 7,747 PICS SNPs that are associated with autoimmune disease or non-immune traits^25^. 4,331 (55.9%) were associated with autoimmune disease, and 3,416 (44.1%) were associated with non-immune traits. In addition we obtained a set of SNPs associated with six overlapping autoimmune diseases using the GRASP catalog (genome-wide significance p-value < 10^-8^).

We constructed a synthetic loop set for immune and non-immune SNPs and any TSS within 1 Mb of each SNP. We then assigned VCsqrt signal in each biological replicate of the three T cell subtypes to the synthetic loop set, as described above.

VCsqrt signal per sample was quantile-quantile normalized as above. In this analysis, we did not restrict to HiCCUPS-identified loops but instead examined all possible interactions between a SNP and TSS within 1 Mb. Many of these interactions do not exist, and therefore had little or no matrix-balanced signal supporting them. While we removed all SNP TSS pairs below an average of 1 normalized read per sample from subsequent analyses, in general these false interactions contributed little to the overall differential signal for a trait.

H3K27ac data was downloaded from the WashU Roadmap repository. PICS SNPs were taken from Farh, et al^25^. Rather than require strict membership within H3K27ac peaks, PICS SNPs were labeled as active if they lied within 8 kb of a peak, raising the number of nominally functional SNPs from ~700 to ~3200 per cell type, out of 7735 total candidate SNPs.

Differential looping across cell types was assessed by one-sided t-test per trait and activity partition if there were at least eight PICS SNP-TSS pairs in the partition. T_H_17 bias was defined as T_H_17 total loop signal minus Naïve total loop signal; Treg bias was defined as Treg total loop signal minus Naïve total loop signal; and Naïve bias was defined as Naïve total loop signal minus one-half times T_H_17 and Treg total loop signal. For the main text figure, Naïve bias was assessed only using SNPs that were active in Naïve cells, T_H_17 bias from SNPs active in T_H_17 and not Naïve, and Treg bias from SNPs active in T_reg_ and not Naïve. p-values were corrected for multiple hypothesis testing by the Holm method using the p.adjust function in R. Bias assessed from SNPs with the opposing cell type specificities (e.g. Naïve bias using SNPs active in T_H_17 and T_reg_ but not Naïve) yielded no significant hits after correction.

### SNP and TSS Connectivity Analysis

Among all immune and non-immune SNPs, 2,562 (33.1%) were located within a 25 kb region centered around TSS, 2,618 (33.8%) were located in a gene body, and the remaining 2,567 (33.1%) were located in intergenic regions.

To capture the confident SNP-TSS connections for SNP-TSS pair identification, we overlapped the anchors of significant loops that were identified by Fit-Hi-C^56^ with the SNP/TSS locations. We identified 14,738 SNP-TSS pairs that were supported by Fit-Hi-C loops, and there were 3,046 unique SNPS connected with at least one gene. Among those 3,046 SNPs, there were 2,181 (71.6%) and 865 (28.4%) SNPs annotated with immune and non-immune SNPs, respectively. As expected, the immune disease SNPs are more likely to connect with genes (Fisher’s exact test, p-value = 4.8 x 10^-85^).

### Phasing of HCASMC Samples

We used BEAGLE 4.1 to impute and phase recalibrated variants using 1000 Genome phase 3 version 5a as a reference panel. The Beagle phasing algorithm was set on the following criteria. At each iteration that the algorithm performs, phased input data are used to build a local model of a haplotype-cluster. After the local haplotype-cluster model is created, for each individual phased haplotypes are sampled using the induced diploid HMM conditional on the individual’s genotypes. The sampled haplotypes are then used as the input to feed in the next iteration of the algorithm. In the final iteration, instead of sampling haplotypes, BEAGLE uses the Viterbi algorithm to select the conditional on the diploid HMM and the individual’s genotype data and to obtain the haplotypes for each individual that possess the greatest probability, and these most-likely haplotypes are the final output of the BEAGLE phasing algorithm.

### Allelic Mapping Bias Simulation

We constructed a personal genome by editing the reference genome (hg19) according to SNP information. SNPs labeled as “1|0” or “1|1” in the vcf file were replaced with the alternative allele for genome 1. SNPs labeled as “0|1” or “1|1” in the vcf file were replaced with the alternative allele for genome 2. Loop anchors were extended 100 bp in both directions and sequences were extracted using samtools for each genome. ~20X reads were simulated for each region, using wgsim with parameters “ -e 0.01 -d 100 -s 20 -1 75 -2 75 -S -1 -h -R 0.1”. Simulated reads were mapped to the “N” masked genome. Mapping parameters were the same used by HiC-Pro (“–very-sensitive -L 30 –score-min L, -0.6, -0.2 –end-to-end –reorder –phred33-quals). Allelic specific reads were separated according to the SNP information and then counted for each loop anchor using bedtools.

### Correction for False Positives

Despite not being originally implemented in enrichment interaction datasets, we have previously demonstrated that KR and VC matrix balancing corrects for false positives caused by high 1D fragment visibility^16^. Loop calls are either from HiCCUPS or Fit Hi-C, which are KR and VC matrix balanced, respectively. For differential analysis of HiCCUPS (Figure 3) and SNP-TSS Fit Hi-C (Figure 4c) loop calls, we are restricted within a set of loops that were obtained from matrix balancing. Therefore, while differential loops can be driven by both changes in looping strength and/or 1D ChIP signal, the final interactions being observed are loops. Additionally, we performed differential analysis of HiCCUPS loop calls using two separate methods – one using

VCsqrt normalized reads and another with non-normalized reads (edgeR) and found high agreement. The SNP-TSS synthetic loop analysis in Figure 4b was not restricted to loop calls, however was performed with VCsqrt matrix balanced reads to avoid false positives. Those SNP-TSS synthetic loops were then subset by overlap with Fit Hi-C for further differential analyses in Figure 4c. Finally, virtual 4C analysis was performed on non-normalized reads to highlight EIS contributions of both 1D and 3D signal changes, however HiCCUPS loop calls are included in the relevant anecdotes.

### ATAC-seq

Cells were isolated and subjected to ATAC-seq as previously described^16^. Briefly, 55,000 cells were pelleted, resuspended in 50 μL lysis buffer (10mM Tris-HCl, pH 7.4, 3mM MgCl2, 10mM NaCl, 0.1% NP-40 (Igepal CA-630), and immediately centrifuged at 500 rcf for 10 min at 4^0^C. The nuclei pellets were resuspended in 50 μL transposition buffer (25 μl 2X TD buffer, 22.5 μL dH_2_0, 2.5 μL Illumina Tn5 transposase), and incubated at 37^0^C for 30 min. Transposed DNA was purified with MinElute PCR Purification Kit (Qiagen), and eluted in 10 μL EB buffer.

### ATAC-seq Data Processing

Adapter sequence trimming using SeqPurge and mapping to hg19 using Bowtie2 were performed. These reads were then filtered for mitochondrial reads, low quality, and PCR duplicates. The filtered reads for each sample were merged and peak calling was performed by MACS2. Each individual sample reads in peaks were quantified using Bedtools intersect with the MACS2 narrow peaks. Peak counts were then combined into a matrix NxM where N represents called peaks and M represents the samples and each value Di,j represents the peak intensity for the respective peak i in sample j. This matrix was then normalized using the “CQN” package in R to minimize bias in GC content and length.

### CRISPRi Validation of HiChIP Targets

For virus production, 5 × 10^6^ of HEK293T cells were plated per 10 cm plate. The following day, plasmid encoding lentivirus was co-transfected with pMD2.G and psPAX2 into the cells using Lipofectamine 3000 (Thermo Fisher, L3000) according to the manufacturer’s instructions. Supernatant containing viral particles was collected 48 hours after transfection and filtered. For lentivirus encoding individual sgRNAs, virus was concentrated 10-fold using Lenti-X concentrator (Clontech, 631232) and stored at - 80°C.

In order to generate a My-La cell line expressing CRISPRi, 2 × 10^6^ of My-La cells were plated per T75 flask. A dCas9-BFP-KRAB-2A-Blast construct was generated by inserting a 2A-Blast cassette into dCas9-BFP-KRAB (Addgene 46911). 24 hours after plating, lentivirus harboring the dCas9-KRAB construct was added with polybrene (4 μg / mL). Media was changed 24 hours after infection, and then again 48 hours after infection with Blasticidin (Thermo Fisher, A1113903) at 4 μg / mL. Blasticidin resistant cells were selected for eight days with changing media every other day.

Three different U6 were used for transcription of three different sgRNAs targeting the candidate enhancers, as previously described^57^. For *MYC* locus CRISPRi experiments, each enhancer was targeted by one guide, and therefore all *MYC* GM or My-La enhancers together were targeted in one experiment. For the PICS SNP CRISPRi experiments, three guides were targeted to a single SNP-containing enhancer. One of three sgRNAs was cloned into a lentiviral vector with a human (pMJ117, Addgene 85997), mouse (pMJ179, Addgene 85996) or bovine (pMJ114, Addgene 85995) U6 promoter. These U6-sgRNA constructs were then combined into lentivirus with a Puromycin-2A-mCherry vector, which was modified from Addgene 46914. My-La-CRISPRi cells were infected with lentivirus harboring 3 sgRNAs and selected by Puromycin (Thermo Fisher, A11138) at a final concentration of 1 μg / mL. Previously reported sgRNAs targeting VPS54 or SEC24C were used for validating CRISPRi functionality in My-La cell line^58^.

For readout of CRISPRi validation, we performed qRT-PCR and cell growth assays on three biological and two technical replicates. For qRT-PCR, RNA was Trizol extracted (Thermo Fisher, 15596026) and purified using the Zymo RNA Clean and Concentrator kit (Zymo Research, R1016). qRT-PCR was performed with Brilliant qRT-PCR mastermix (Agilent, 600825). Ct values were measured by using Lightcycler 480 (Roche) and relative expression level was calculated by ddCt method compared to a *GAPDH* control. Primer sequences are listed in **Supplementary Table 6**. For cell growth, we used the CellTiter-Glo kit (Promega, G7572) according to the manufacturers instructions. Statistics for both RNA and cell growth changes were calculated using a Student’s t test against the non-targeting control.

### CRISPRa Validation of HiChIP Targets

Jurkat cells were transduced with a lentiviral dCas9-VP64-2A-GFP expression vector (Addgene 61422). Single GFP+ cells were sorted by FACS into the wells of a 96-well plate, and a clone with bright uniform GFP expression were selected for use in future experiments.

sgRNAs were cloned in arrayed format for CD69 HiChIP peaks falling outside the range of the tiling CRISPRa screen^26^. sgRNAs were chosen based on high predicted on-target activity^59^ and low predicted off-target activity^60^. sgRNAs were cloned into the lentiviral expression vector “pCRISPRia-v2” (Addgene 84832) as described in Horlbeck et al^61^. Lentivirus was produced by transfecting HEK293T with standard packaging vectors using *TransIT-LTI* Transfection Reagent (Mirus, MIR 2306). Media was changed 24 hours post-transfection. Viral supernatant was harvested at 48 and 72 hr following transfection and immediately used for infection of Jurkat-dCas9-VP64 cells.

Jurkat-dCas9-VP64 cells were infected with lentiviral sgRNAs by resuspending cells in a 1:1 mix of fresh media and lentiviral supernatant at a final concentration of 0.25 × 10^6^ cells/mL with 5 μg / mL polybrene. Cells were spinfected for 1 hour at 1000 rcf, 32 °C.

The next day, half of the media was removed and replaced with fresh lentiviral supernatant, and the spinfection was repeated. The next day, the cells were resuspended in fresh media with 1.5 μg / mL puromycin and cultured for 2 days to remove uninfected cells. For readout of CRISPRa validation, we performed qRT-PCR and FACS on two biological and two technical replicates. RNA extraction and qRT-PCR was performed as described above. Expression of CD69 on infected cells (GFP+BFP+) was analyzed by flow cytometry with an Attune NxT flow cytometer (Life Technologies). Statistics for both RNA and protein level changes were calculated with a one-way ANOVA followed by a Dunnet’s multiple comparisons test against the non-targeting control.

---

**Supplementary Figure 1. H3K27ac HiChIP enriches E-P-associated chromatin contacts. (A)** Schematic of chromatin contacts captured in H3K27ac HiChIP. **(B)** Loop call overlap for cohesin HiChIP and H3K27ac HiChIP in mES cells. **(C)** Contact distance distribution for loops that are biased in cohesin versus H3K27ac HiChIP. **(D)** Proportion of cohesin and H3K27ac biased HiChIP loops that have cohesin, CTCF, H3K27ac, and RNA polymerase II binding in at least one loop anchor. **(E)** Virtual 4C interaction profile of an H3K27ac-biased loop focused at the *MALAT1* promoter. **(F)** Virtual 4C interaction profile of a cohesin-biased loop with associated low transcriptional activity.

**Supplementary Figure 2. H3K27ac HiChIP achieves high chromatin loop signal-over-background at low cell inputs. (A)** KR balanced interaction matrices focused around the *Etv5* locus in mES cells with decreasing cellular starting material. **(B)** Read support reproducibility of loops in H3K27ac HiChIP libraries from 25 million cells compared to HiChIP in lower cell input libraries. **(C)** Aggregate peak analysis of loops in mES H3K27ac HiChIP libraries.

**Supplementary Figure 3. H3K27ac HiChIP generates reproducible chromatin loop signals at low cell inputs. (A)** Comparison of KR balanced interaction maps in H3K27ac HiChIP biological replicates. **(B)** Read support reproducibility of loops between H3K27ac HiChIP biological replicates. **(C)** HiCCUPS loop call overlap between H3K27ac HiChIP libraries from 25 million and 50 thousand mES cells. **(D)** Preseq library complexity estimation of H3K27ac HiChIP libraries from 25 million and 50 thousand mES cells.

**Supplementary Figure 4. H3K27ac HiChIP biological replicates from primary sorted T cells are highly reproducible. (A)** FACS strategy for Naïve, T_H_17, and Treg cells starting from human peripheral blood. Post-sort validation was used to ensure purity of T cell subtypes. Number represents percent of cells within that gate. **(B)** KR balanced interaction map of T cell subtype biological replicates. **(C)** Read support reproducibility of loops between H3K27ac HiChIP biological replicates in Naïve, T_H_17, and Treg cells. **(D)** Aggregate peak analysis of loops in Naïve, T_H_17, and Treg H3K27ac HiChIP libraries.

**Supplementary Figure 5. Validation of HiChIP-identified *CD69* distal enhancers with CRISPR activation**. CRISPRa validation in Jurkat cells of *CD69* distal enhancers. *CD69* protein levels are shown for individual sgRNAs tiling H3K27ac HiChIP-identified distal *CD69* enhancers relative to the *KLRF2* promoter as a locus negative control and a non-targeting negative control.

**Supplementary Figure 6. Global enhancer connectome characterization in T cell differentiation. (A)** ChromHMM classification of union T cell loop anchors. **(B)** Contact distance distribution of union T cell loops. **(C)** Agreement in residual signal observed between sample signals per loop after removing imputed shared signal, calibrated by a null distribution of random pairings of loops. QQ plot shows modest enrichment above random pairings. PCA on residual signal clusters samples first by Naive vs Memory cell types (PC1), and then by donor identity (PC2, 3). **(D)** Overlap of differential interactions between Naïve, T_H_17, and Treg subtypes. Biased interactions were obtained by performing pairwise comparisons between T cell types and analyzing the top 5% enriched and top 5% depleted EIS in each T cell subtype. **(E)** ChromHMM annotation of total loops, differential loops, and shared loops in all three T cell subtypes. O corresponds to other loop anchors not classified as enhancer or promoter. **(F)** Number of connections for different classes of loop elements. **(G)** Quantification of promoters skipped before an enhancer reaches its loop target.

**Supplementary Figure 7. Positioning of differential HiChIP contacts in gene dense chromosomes. (A)** Distribution of T cell subtype differential HiChIP contacts across different chromosomes compared to the distribution of all loops. **(B)** Correlation of differential loop density with gene density and GC content.

**Supplementary Figure 8. Characterization of conformational landscapes surrounding key T cell regulatory factors. (A – C)** Virtual 4C interaction profiles anchored at the promoters of canonical Naïve, T_H_17, and Treg regulatory factors.

**Supplementary Figure 9. Chromosome conformation dynamics of canonical T cell regulatory factors. (A – C)** Delta interaction maps focused around known Naïve, T_H_17, and Treg regulatory factors.

**Supplementary Figure 10. Contribution of H3K27ac ChIP and chromosome conformation to HiChIP EIS. (A)** (Left) Proportion of H3K27ac ChIP peaks within EIS differential loop anchors that are cell-type specific (log_2_ fold change > 1) or shared across all three subtypes. (Right) Global correlation of EIS and H3K27ac ChIP fold-change in different T cell subset pairwise comparisons. **(B)** Same as (A), but using HiChIP 1D differential signal at EIS biased loop anchors. **(C)** Overlap of H3K27ac HiChIP and bins of Hi-C loops with increasing T cell subset and GM H3K27ac ChIP-seq signal. **(D)** Overlap of CD4+ Capture Hi-C^14^ with total and differential T cell subset HiChIP loops. **(E)** T_reg_-specific loops at the *LRRC32* promoter not observed in other H3K27ac HiChIP T cell subsets nor in CD4+ Capture Hi-C data^14^.

**Supplementary Figure 11. Enrichment of autoimmune SNPs in T cell HiChIP loop anchors. (A)** Enrichment of specific PICS autoimmune disease and non-immune SNPs in anchors of loops called by Juicer and Fit-Hi-C compared to a background shuffled loop set. **(B)** Enrichment of all PICS autoimmune disease and non-immune SNPs in T cell subset biased loop anchors and all anchors compared to a background shuffled loop set.

**Supplementary Figure 12. T cell subtype HiChIP specificity of autoimmune SNPs. (A)** H3K27ac HiChIP signal bias in T cell subsets for PICS SNP-TSS pairs grouped by each SNP’s presence in cell type-specific or shared H3K27ac ChIP peaks up to 8 kb away. **(B)** H3K27ac HiChIP signal bias in T cell subsets for PICS SNP-TSS pairs grouped by each SNP’s presence in cell type-specific or shared H3K27ac ChIP peaks up to 2.5 kb away. **(C)** H3K27ac HiChIP signal bias in GM, K562, and My-La cell lines for PICS SNP-TSS pairs grouped by each SNP’s presence in T cell subset-specific or shared H3K27ac ChIP peaks up to 8 kb away. **(D)** H3K27ac HiChIP signal bias in T cell subsets for GRASP SNP-TSS pairs grouped by each SNP’s presence in cell type-specific or shared H3K27ac ChIP peaks up to 2.5 kb away. **(E)** Average number of HiChIP gene targets for non-genic autoimmune disease and non-immune SNPs. **(F)** Quantification of promoters skipped before a SNP reaches its gene target. **(G)** Quantification of SNP HiChIP gene targets in autoimmune disease.

**Supplementary Figure 13. Validation of HiChIP signal at SNP-eQTL contacts. (A)** Validation of HiChIP signal at SNP-TSS pairs using interaction profiles of eQTL SNPs to ensure they contact their associated target gene promoter. **(B)** Interaction profiles of CRISPRi-validated loci in My-La.

**Supplementary Figure 14. H3K27ac HiChIP fine-mapping of GWAS variants in haplotype blocks. (A)** Global validation of HiChIP signal at putatively causal SNPs versus corresponding SNPs in LD (r^2^ ≥ 0.8) for Naïve and Treg cell subtypes. SNP-TSS pairs were generated from published fine-mapped datasets, compared to a distance-matched SNP-TSS pair set in the same LD block. **(B)** Interaction profile of the *SATB1* promoter, and a 1 kb resolution visualization of the SNP-containing enhancer of interest. Changes in bias between 5 kb and 1 kb resolution reflects EIS focused 1 kb around the SATB1 TSS and specific SNPs within the enhancer. LD SNPs (r^2^ ≥ 0.8) correspond to GRASP SNPs (genome-wide significance p-value < 10^-8^). The highlighted SNP is a PICS closest to focal EIS to *SATB1*.

**Supplementary Figure 15. Chromatin interaction landscape of the 9p21.3 cardiovascular disease risk locus**. HCASMC v4C interaction profiles focused around the promoters of *CDKN2A, CDKN2B*, and *ANRIL* within the 9p21.3 locus.

**Supplementary Table 1. HiChIP data processing metrics**. HiC-Pro mapping, Hi-C filtering, duplicate removal, and interaction length statistics for all HiChIP libraries.

**Supplementary Table 2. HiCCUPS high confidence loop calls**. High confidence loop calls for all HiChIP libraries.

**Supplementary Table 3. HiCCUPS differential EIS in T cell subtypes by edgeR**. High confidence loop calls in T cell subsets with edgeR significance and fold-change for pair-wise comparisons.

**Supplementary Table 4. HiCCUPS differential EIS in T cell subtypes**. Top 5% of EIS ranked by cell-type bias for each pair-wise comparison between different T cell subtypes.

**Supplementary Table 5. HiChIP gene targets of autoimmune and CAD SNPs**. SNP type (within gene, at TSS, or intergenic), associated disease, and corresponding loop annotated with the HiChIP gene target. Included are edgeR significance and fold-change for cell-type comparisons of interest.

**Supplementary Table 6. sgRNA and primer oligonucleotide sequences**. sgRNA and primer sequences used throughout the study.

## DATA AVAILABILITY STATEMENT

Raw and processed data available at NCBI Gene Expression Omnibus, accession number GSE101498.

T cell ATAC and HiChIP datasets can be visualized in the WashU Epigenome Browser with the following link:

http://epigenomegateway.wustl.edu/browser/?genome=hg19&session=YAIzYBfrl9&statusld=1698051079

